# Genomic Decoding of Specialized Aromatic Hydrocarbon Degradation in Mangrove-Derived *Gordonia* sp. B7-2

**DOI:** 10.64898/2026.05.23.727409

**Authors:** Fangyan Jiang, Haodong Shi, Meilin Lu, Zhenqi Zhao, Xiaoxiong Xu, Huimin Feng

**Author notes:** **Correspondence:** Huimin Feng, Xiaoxiong Xu. These authors contributed equally to this work (Fangyan Jiang, Haodong Shi).

## Abstract

Petroleum pollution has increased worldwide, driving the search for microorganisms with efficient hydrocarbon-degrading capabilities. Here, we report a novel bacterium, *Gordonia* sp. B7-2, isolated from mangrove sediments in Hainan, China. Phylogenetic analysis based on the 16S rRNA gene and whole-genome sequences, together with digital DNA–DNA hybridization and average nucleotide identity values, supported its classification as a new species within the genus *Gordonia*. The complete genome of strain B7-2 consists of a single circular chromosome of 5.39 Mb with a G+C content of 65.99%, and encodes 4,887 protein-coding genes. Genomic annotation revealed a complete pathway for aromatic hydrocarbon degradation, including genes encoding protocatechuate 3,4-dioxygenase and biphenyl-2,3-diol 1,2-dioxygenase, whereas genes involved in the initial oxidation of alkanes were absent. Consistent with these genomic predictions, strain B7-2 degraded 64.33% of crude oil (300 mg/L) within 28 days, with rapid degradation during the initial 14 days, followed by a slower phase thereafter, reflecting the dynamics of complex hydrocarbon mixtures. Together, these results demonstrate that strain B7-2 is specialized for the degradation of aromatic hydrocarbons and highlight its potential for targeted petroleum bioremediation.

**IMPORTANCE:** Mangrove ecosystems are highly vulnerable to petroleum contamination, yet the microorganisms responsible for hydrocarbon turnover in these environments remain poorly characterized. This study describes a new bacterial species, Gordonia sp. B7-2, that exhibits a strong metabolic specialization for aromatic hydrocarbon degradation. Unlike many known oil-degrading bacteria that preferentially utilize alkanes, strain B7-2 targets aromatic components of crude oil, which are among the most persistent and toxic fractions.Its ability to efficiently degrade crude oil highlights its potential in the bioremediation of contaminated coastal environments and expands our understanding of microbial contributions to hydrocarbon cycling in mangrove sediments

## Background

Petroleum, commonly referred to as crude oil, is a naturally occurring liquid hydrocarbon system composed of a complex mixture of organic compounds. Its constituents typically include saturated hydrocarbons (e.g., alkanes), aromatic hydrocarbons, and polar components such as asphaltenes and resins (1–3). Over the past two decades, rapid industrialization has driven a marked increase in the production and consumption of petroleum products (4). Global crude oil demand is projected to rise from 84 million barrels per day in 2000 to 102 million barrels per day by 2023 (5). Petroleum supply chain operations are frequently associated with unintended hydrocarbon spills into ecosystems due to anthropogenic factors (6). Oil spills pose a serious threat to ecosystems (7). Consequently, the removal of excessive petroleum hydrocarbons from contaminated environments has become an urgent environmental challenge.

The development of petroleum-degradation technologies remains an active and expanding field of research (5). Various physical and chemical approaches, including photodegradation, incineration, adsorption, separation, thermal washing, and *in situ* chemical oxidation, have been used for petroleum remediation (8–10). However, conventional physicochemical methods often generate ancillary contaminants, leading to additional environmental contamination, and suffer from inherent limitations such as low remediation efficiency and incomplete resource recovery (11). As an environmentally sustainable alternative, bioremediation has emerged as a promising strategy for the degradation of petroleum hydrocarbons. This approach harnesses microbial metabolic capabilities, utilizing bacteria/yeasts (bacterial remediation), algae (phycoremediation), and fungi (mycoremediation) to enzymatically degrade petroleum contaminants through natural biodegradation pathways (7). In recent years, numerous petroleum-degrading microorganisms isolated from soil and other environments have been identified, including species belonging to the genera *Aspergillus*, *Penicillium*, *Fusarium*, *Trametes*, *Trichoderma* (12–14), *Bacillus*, *Pseudomonas*, *Acinetobacter*, *Stenotrophomonas*, *Serratia*, *Rhodococcus*, and *Gordonia* (15–20).

Among these, the genus *Gordonia* has been investigated for its remarkable capacity to degrade a wide range of hydrocarbons, including petroleum, diesel, aliphatic and aromatic hydrocarbons, polycyclic aromatic hydrocarbons (PAHs), and phthalates. Yang *et al.* reported that an immobilized bacterial agent of *G. alkanivorans* W33, developed through high-density fermentation combined with bacterial immobilization technology, achieved 56.3% petroleum degradation (250.2 mg/kg/d) in soil containing 20,000 mg/kg hydrocarbons over 45 days (19). *G. amicalis* HS-11 exhibited a diesel oil degradation efficiency of 92.85 ± 3.42% (1% [v/v]) after 16 days of aerobic incubation (21). Strain *G. iterans* Co17 was shown to utilize anthracene and naphthalene in oil, with degradation rates of 63.2% and 55.3%, respectively (22). Frantsuzova *et al.* (23) observed the ability of *G. polyisoprenivorans* 135 to degrade a wide range of PAHs. Hu *et al.* (24) reported that *Gordonia* sp. GZ-YC7 degraded 45% of di-(2-ethylhexyl) phthalate (DEHP) at a high concentration of 500 mg/kg in soil within 5 days and exhibited exceptional tolerance, with optimal degradation at 1,000 mg/L and sustained activity at concentrations up to 4,000 mg/L in liquid culture. In addition, *G*. *polyisoprenivorans* B251 exhibited polyethylene-degrading capacity, as evidenced by characteristic chemical modifications, including changes in the carbonyl index, and surface morphological alterations following a 30-day polyethylene film incubation (25). These inherent metabolic adaptabilities highlight *Gordonia* spp. as promising candidates for petroleum hydrocarbon bioremediation and broader environmental biotechnological applications.

Despite the growing number of petroleum-degrading strains reported within the genus *Gordonia*, genomic and genetic information for many of these strains remains limited, thereby constraining research on the molecular mechanisms underlying petroleum degradation. In this study, we report a novel *Gordonia* strain, designated B7-2, isolated from mangrove sediments in Hainan, China. Strain B7-2 is a Gram-positive, aerobic, and non-motile bacterium. Its complete genome was sequenced, and putative proteins were annotated. Phylogenetic analysis indicated that strain B7-2 represents a new species within the genus *Gordonia*. Genes associated with petroleum degradation were predicted using antiSMASH and homologous BLAST analyses. Furthermore, comparative genomic analyses were conducted to elucidate the phylogenetic relationships and metabolic potential of strain B7-2 in relation to other *Gordonia* species. This study provides valuable insights into the genomic characteristics of petroleum-degrading bacteria, thereby contributing to a deeper understanding of the molecular mechanisms involved in petroleum hydrocarbon degradation and supporting the development of targeted bioremediation strategies.

## MATERIALS AND METHODS

### Bacterial Strains

The strain *Gordonia* sp. B7-2 was isolated from mangrove sediments collected at Qingmei Port, Sanya, Hainan, China (18°13′50.9″N 109°37′15.9″E), in June 2023 using a standard dilution plating method. *Gordonia* sp. B7-2 was deposited at the China Center for Type Culture Collection (CCTCC), China, under the number CCTCCAA2024105.

### Culture Media

ISP2 liquid medium was prepared by dissolving yeast extract (4 g), malt extract (10 g), and glucose (4 g) in 1 L of distilled water, followed by pH adjustment to 7.2–7.4 and sterilization at 121 °C for 20 min. The corresponding solid medium was prepared by supplementing the same formulation with 20 g/L agar before sterilization.

For crude oil biodegradation assays, a basal salts medium was used, consisting of (per liter of distilled water): NaCl (29 g), Na₂HPO₄ (3 g), KH₂PO₄ (1 g), NaH₂PO₄ (1 g), KNO₃ (1 g), NH₄Cl (1 g), MgSO₄·7H₂O (0.7 g), and 1 mL of a trace element solution. The trace element solution contained CaCl₂ (20 mg/L), FeCl₃ (30 mg/L), CuSO₄ (0.5 mg/L), MnSO₄·H₂O (0.5 mg/L), and ZnSO₄·7H₂O (10 mg/L). The basal salts medium was sterilized at 121 °C for 20 min. The final crude oil medium was prepared by adding 300 mg of ether-sterilized crude oil to 1 L of sterile basal salts medium.

### Strain Cultivation

The strain was inoculated into ISP2 liquid medium and cultured in a shaking flask at 28 °C and 180 rpm for 7 days. Cells were then harvested by centrifugation, washed three times with sterile buffer, and resuspended in ISP2 medium containing 20% (v/v) glycerol for preservation at −80 °C until further use.

### Genomic DNA Extraction

*Gordonia* sp. B7-2 was streaked onto ISP2 solid medium and incubated at 28 ℃ for 7 days. A single colony was then inoculated into 50 mL of ISP2 liquid medium and cultured at 28 ℃ and 180 rpm for approximately 5 days to prepare the seed culture. Subsequently, 2 mL of the seed culture was transferred into 200 mL of fresh ISP2 liquid medium for secondary cultivation under identical conditions (28 ℃, 180 rpm) for an additional 7 days. After cultivation, the cell biomass was harvested by centrifugation at 7,000 × *g* for 10 min at 4 ℃ and washed with sterile water to obtain fresh cell pellets. Genomic DNA was extracted using a Bacterial Genomic DNA Extraction kit (magnetic beads method) (Tiangen Co., Ltd., Beijing, China) according to the manufacturer’s protocol. The extracted DNA was purified and quantified, and high-quality DNA was selected for subsequent analyses.

### Genome Sequencing and Assembly

Whole-genome sequencing of strain B7-2 was performed using both the PacBio Sequel II (PacBio, USA) and the Illumina HiSeq 2000 platform (Illumina, USA) at Majorbio Bio-pharm Technology Co., Ltd. (Shanghai, China). The combined sequencing data yielded a minimum coverage of 100× to ensure high-quality genome assembly. Genome assembly was performed conducted following methods described previously (26, 27).

### Phylogenetic Analysis

Phylogenetic analyses were conducted based on 16S rRNA gene sequences and whole-genome data. The 16S rRNA gene the sequence of *Gordonia* sp. B7-2 was compared with those of related type strains using BLAST on the NCBI platform. A neighbor-joining phylogenetic tree was constructed using MEGA version 11.0 (28), with *Streptomyces griseoincarnatus* LMG 19316ᵀ (AJ781321) serving as the outgroup. Whole-genome phylogenetic analysis was conducted using the Type Strain Genome Server (TYGS) platform (https://tygs.dsmz.de/). Corresponding average nucleotide identity (ANI) and digital DNA–DNA hybridization (dDDH) values were calculated to evaluate genomic relatedness.

### Genome Annotation

The genome of strain B7-2 was annotated using the Majorbio Cloud Platform. For structural annotation, coding sequences (CDSs) were predicted using Prodigal (29). tRNA and rRNA genes were identified using tRNAscan-SE (30) and Barrnap, respectively. Potential sRNAs were annotated using the Infernal software in conjunction with the Rfam database. Tandem repeats and known repetitive sequences were detected using Tandem Repeats Finder and RepeatMasker. Additional genomic features were predicted using IslandPath-DIMOB and Islander for genomic islands, Phage_Finder for prophages, ISEScan for insertion sequences, TransposonPSI for transposons, and Minced for potential CRISPR-Cas systems.

For functional annotation, the predicted CDSs were aligned against the non-redundant (NR), Swiss-Prot, Pfam, GO, COG, KEGG, and CAZy databases using Diamond, Blast2GO, and HMMER3. The best-matching hits (E-value<1×10⁻⁵) were assigned to each query sequence. Secondary metabolite biosynthetic gene clusters (BGCs) were identified using antiSMASH version 6.0 (31).

### Petroleum Degradation by *Gordonia sp.* B7-2

The petroleum degradation capability of *Gordonia* sp. B7-2 was assessed in minimal salts medium containing crude oil (300 mg/L) as the sole carbon source. Seed cultures were prepared as described in section 2.3, with the modification that harvested cells were washed twice with sterile water and resuspended to an OD_600_ of 1.0. A 2% (v/v) aliquot of this suspension was used to inoculate the crude oil medium. A control group supplemented with an equal volume of sterile ISP2 liquid medium instead of the bacterial suspension was included to account for abiotic losses. All cultures were incubated in triplicate at 28 °C with shaking at 180 rpm for 28 days. Residual crude oil concentrations were measured spectrophotometrically after 7, 14, 21, and 28 days of cultivation.

### Determination of Petroleum Content

Petroleum content was determined as described by Yang *et al.* (19), with some modifications. Petroleum standard solutions (5,000 mg/L) were prepared using petroleum ether as the solvent and serially diluted to obtain a calibration range of 0–500 mg/L. A full scan of the petroleum-petroleum ether standard solution was performed to obtain the appropriate wavelengths. Petroleum concentrations were subsequently measured using a UV-visible spectrophotometer at a maximum absorbance wavelength of 600 nm. A standard curve was constructed by plotting absorbance against petroleum concentration.

After the petroleum degradation experiments, cell biomass was removed from the culture medium by centrifugation at 9,000 × *g* for 20 min. Residual crude petroleum in the supernatant was extracted with petroleum ether using a separatory funnel. The extract was diluted to a final volume of 100 mL in a volumetric flask, and 3 mL of the diluted extract was used to measure absorbance (OD_600_) in quartz cuvettes. Residual petroleum concentrations were calculated using the standard curve. All assays were performed in triplicate, and mean values were used for subsequent calculations.

Crude petroleum degradation efficiency (%) was calculated using the following formula:

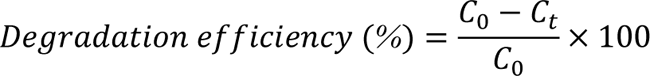

where *C*_0_ is the initial petroleum concentration (mg/L) and *C_t_* is the residual petroleum concentration (mg/L) at time *t*.

## RESULTS

### General Characteristics of the B7-2 Genome

The genome of *Gordonia* sp. B7-2 was assembled into a single circular chromosome of 5,393,941 bp with a G+C content of 65.99% (Table S1). Genome annotation revealed 4,887 protein-coding genes, collectively accounting for 4,879,074 bp (90.45%) of the genome with an average gene length of 998.38 bp. In addition, the genome harbored 218 tandem repeat regions (0.44% of the genome), along with genes encoding 46 tRNAs, 6 rRNAs (two copies each of 5S, 16S, and 23S), and 39 sRNAs. Furthermore, 20 genomic islands and two prophage regions were identified, spanning a total of 507,403 bp. A circular genome map was generated using Circos software (Figure 1). The complete genome sequence has been deposited in the NCBI database under the accession number CP194566 (https://www.ncbi.nlm.nih.gov/nuccore/CP194566.1).

**FIG 1.**
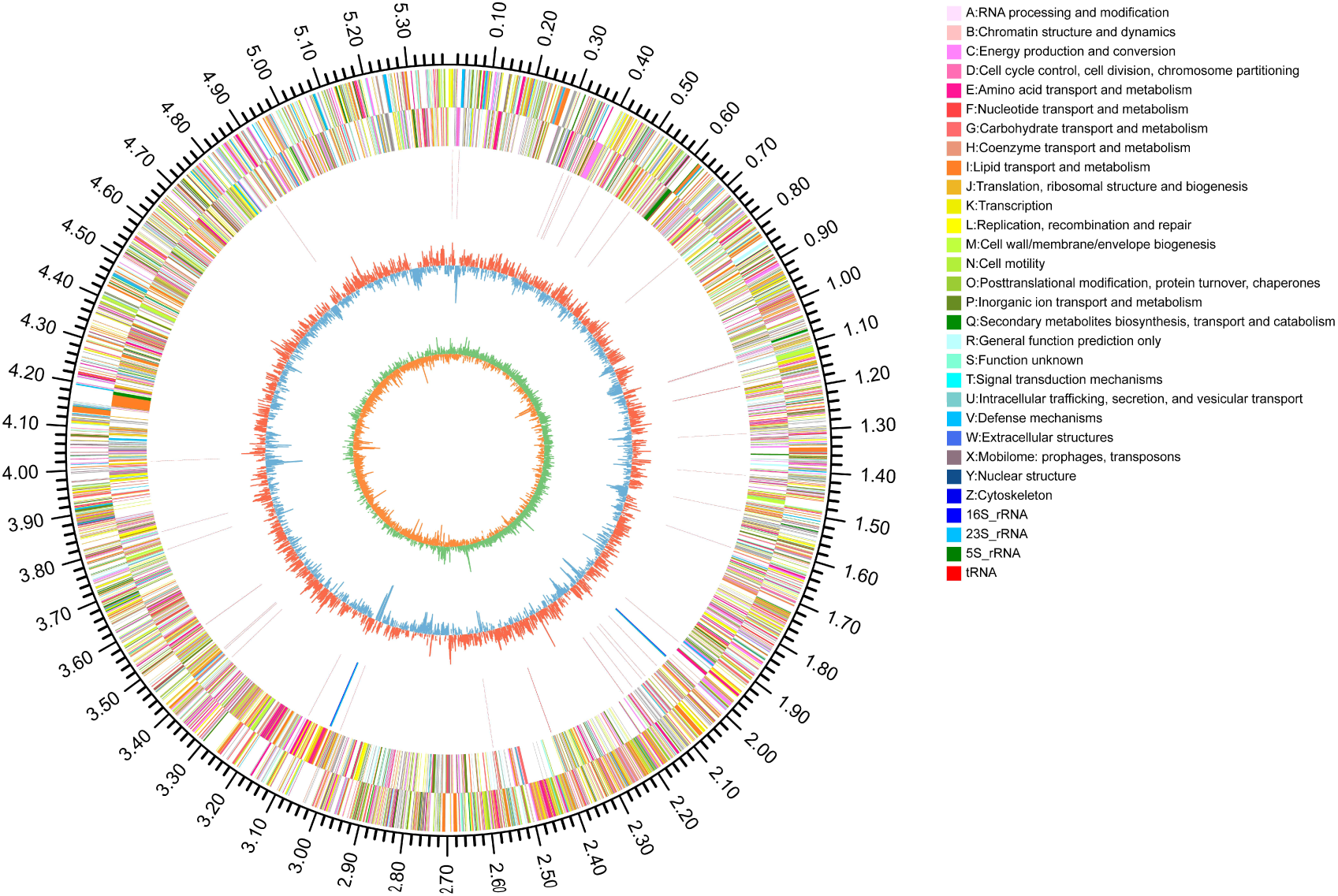
Circular genome map of *Gordonia* sp. B7-2. From the outside in, the first circle shows the genome scale; the second and third circles represent coding sequences (CDSs) on the forward and reverse strands, respectively, colored according to COG functional categories; the fourth circle displays rRNA (red) and tRNA (blue) genes; the fifth circle indicates GC content, with outward red and inward blue peaks representing regions above and below the genomic average, respectively; the innermost circle depicts GC skew (G−C)/(G+C), which is useful for distinguishing leading and lagging strands and identifying replication origins and termini.

### Phylogenetic Analysis of Strain B7-2

The phylogenetic position of strain B7-2 was initially evaluated based on 16S rRNA gene sequence analysis. In the neighbor-joining tree (Figure 2), strain B7-2 formed a robust clade with *Gordonia polyisoprenivorans* NBRC 16320ᵀ (98.52% similarity) and *Gordonia oryzae* RS15-1Sᵀ (98.15% similarity). The maximum 16S rRNA gene sequence similarity between strain B7-2 and its closest relatives was below 99%, suggesting that this strain may represent a novel species within the genus *Gordonia*.

**FIG 2.**
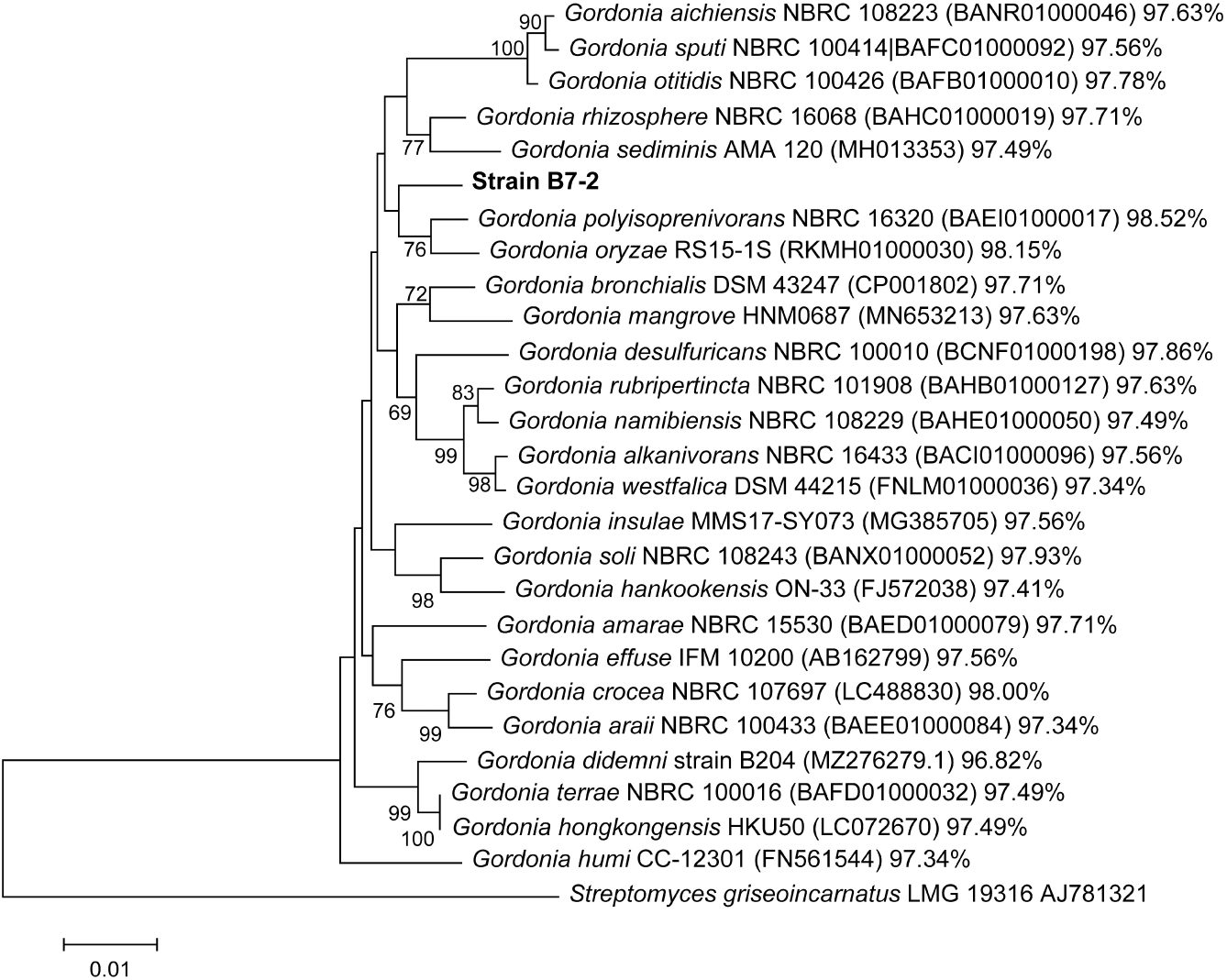
Neighbor-joining phylogenetic tree of *Gordonia* sp. B7-2 and related taxa based on 16S rRNA gene sequences. Bootstrap values (1000 replicates) greater than 50% are shown at branch nodes. The tree was constructed using MEGA version 11.0. The scale bar indicates the number of base substitutions per site.

To further clarify its taxonomic status, whole-genome sequence-based analyses were performed. In the phylogenomic tree (Figure 3), strain B7-2 formed a distinct branch separate from other known *Gordonia* species. The ANI values between strain B7-2 and its closest relatives ranged from 74.31% to 77.63%, while the dDDH values ranged from 20.1% to 23.0%. These values were well below the accepted thresholds for species demarcation (ANI < 95–96%; dDDH < 70%). Notably, the type strains *G. polyisoprenivorans* NBRC 16320ᵀ and *G. oryzae* RS15-1Sᵀ, which showed the highest 16S rRNA similarities, exhibited ANI/dDDH values of 75.90%/21.2% and 75.83%/20.7%, respectively, when compared with strain B7-2. Collectively, these genomic analyses strongly support the conclusion that strain B7-2 represents a novel species within the genus *Gordonia*.

**FIG 3.**
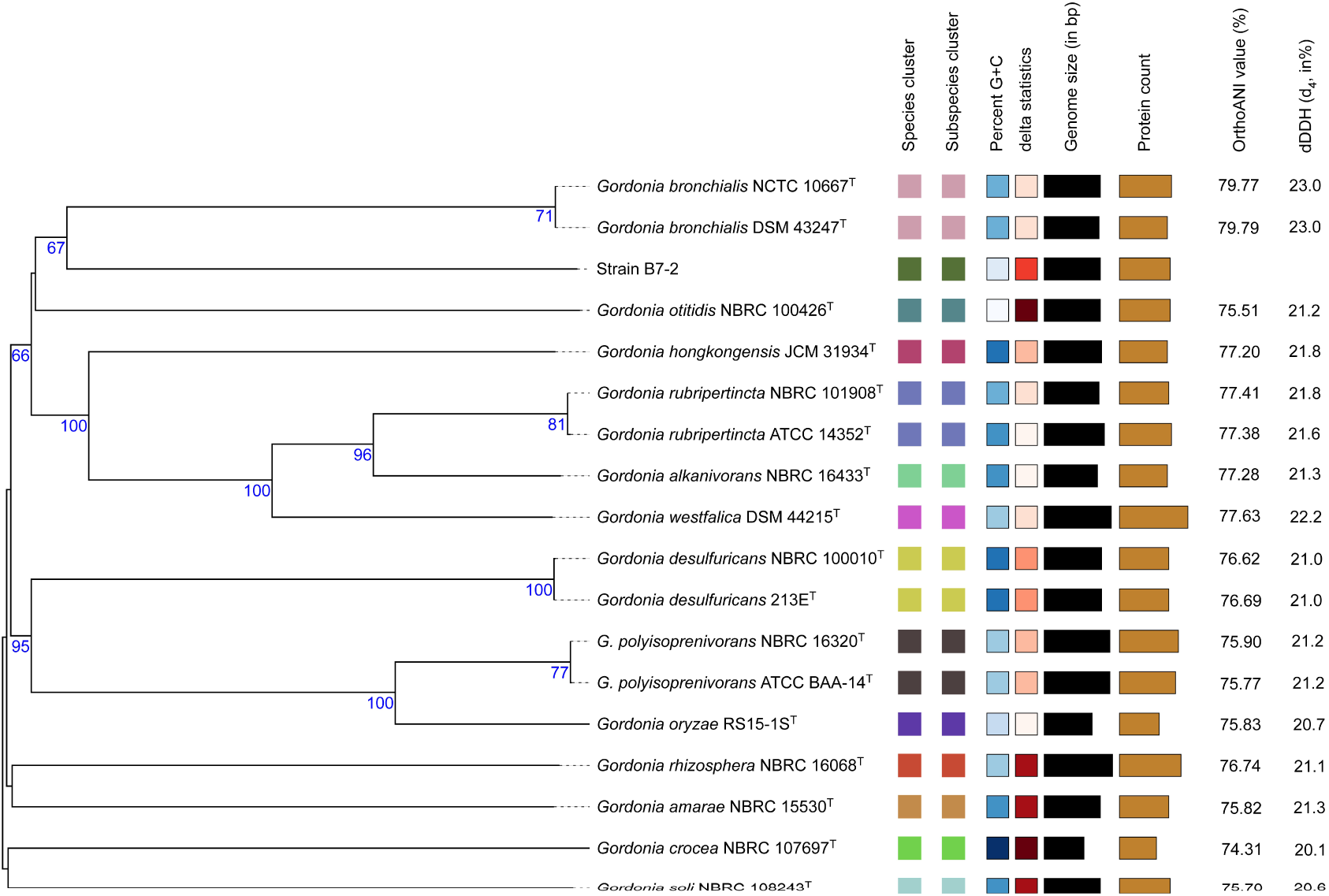
Phylogenomic tree of *Gordonia* sp. B7-2 and related type strains reconstructed from whole-genome sequences using the TYGS platform. Average nucleotide identity (ANI) and digital DNA–DNA hybridization (dDDH) values between strains are shown. T, Type strain.

### Functional Annotation of *Gordonia* sp. B7-2

A total of 4,887 CDSs were annotated in the genome of *Gordonia* sp. B7-2, with a combined length of 4,879,074 bp, accounting for 90.45% of the total genome (Table 1). Among these, 3,313 genes were assigned GO terms, 3,821 genes were annotated in the COG database, 2,240 genes were linked to KEGG pathways, 4,813 genes were matched in the NR database, and 3,400 genes were annotated in the Swiss-Prot database.

**TABLE 1.**
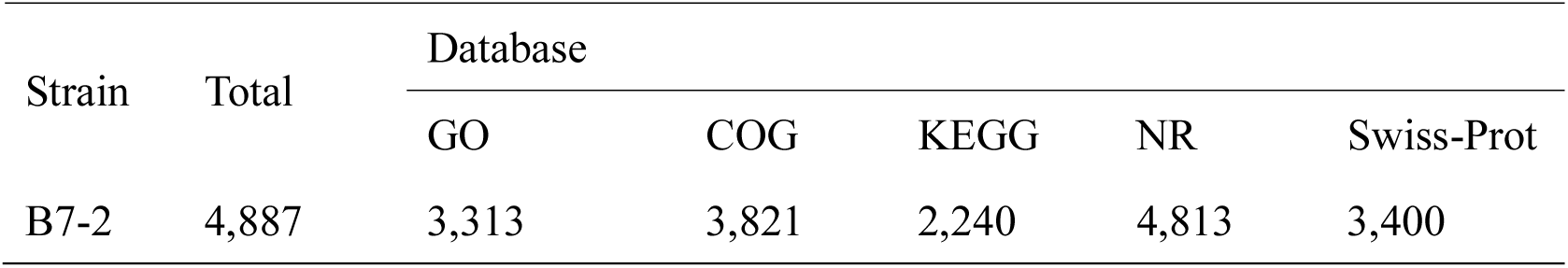
Summary of functional annotation for the protein-coding genes of *Gordonia* sp. B7-2.

### GO Database Annotation

GO classification assigned functional annotations to 3,313 genes (67.79% of all CDSs), which were distributed across three major categories: biological processes (BP), cellular component (CC), and molecular function (MF). As shown in Fig. 4, these genes were distributed across 42 functional subcategories. Within the BP category, 1,494 genes were annotated, with the most abundant functions being the regulation of DNA-templated transcription, translation, and phosphorylation. In the CC category, 1,250 genes were annotated, predominantly associated with integral components of the membrane, cytoplasm, and plasma membrane. The MF category contained 2,748 genes, with DNA binding, ATP binding, and metal ion binding representing the three most prevalent functions.

**FIG 4.**
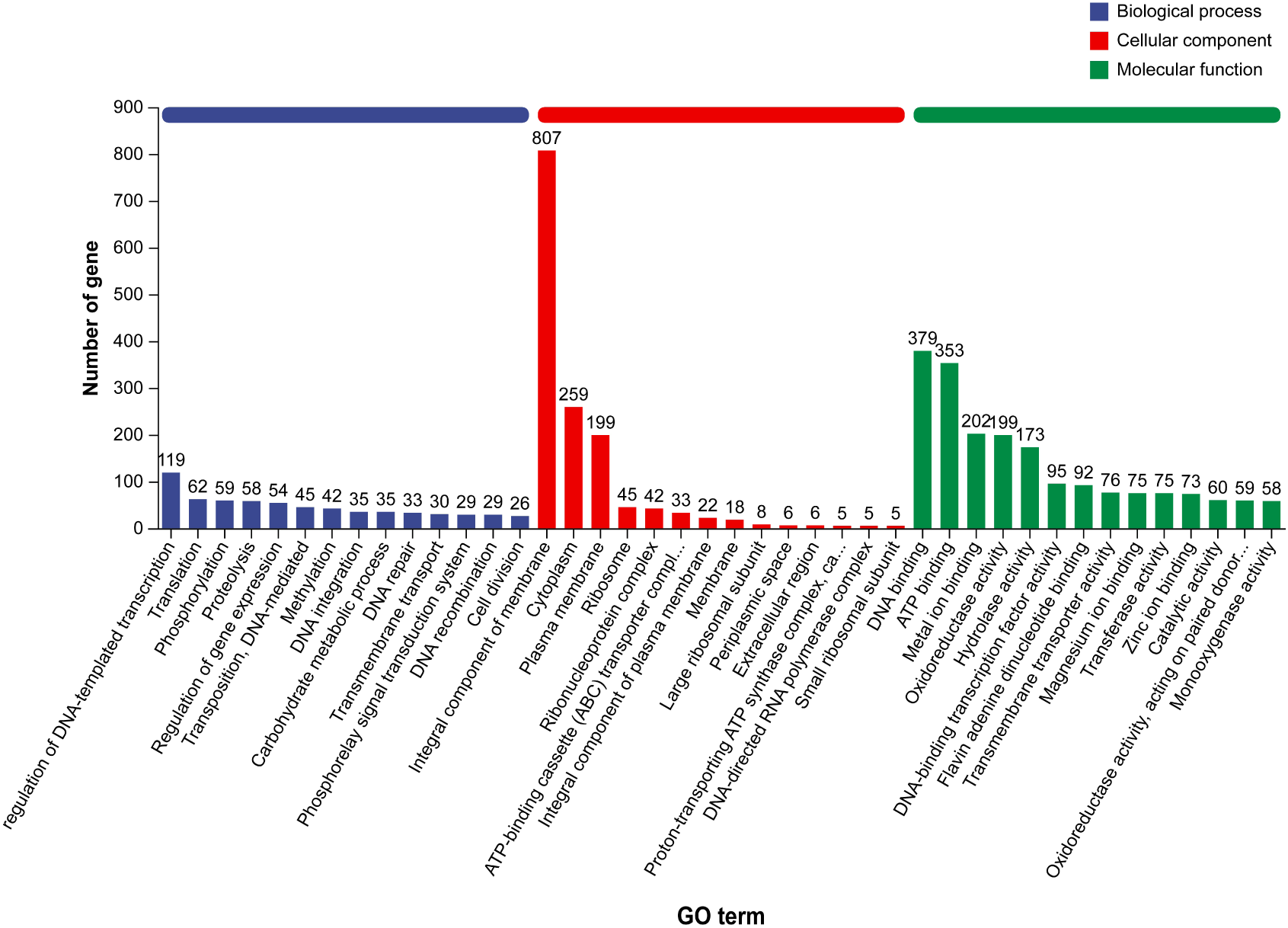
GO classification of the *Gordonia* sp. B7-2 genome.

Based on GO annotations associated with aromatic compound catabolism, alkanal oxidation, fatty acid metabolism, and related enzymatic activities, including dioxygenases, aldolases, and acyltransferases, several key genes implicated in petroleum hydrocarbon degradation were identified. These included protocatechuate 3,4-dioxygenase subunit alpha/beta (gene4336, gene4337), which function as central enzymes in aromatic hydrocarbon degradation; biphenyl-2,3-diol 1,2-dioxygenase (gene0804), which is involved in the breakdown of polycyclic aromatic hydrocarbons; and 4-hydroxy-2-oxovalerate aldolase (gene0791, gene2627, and gene4366), a terminal metabolic enzyme in the aromatic hydrocarbon pathway. In addition, FMN-linked alkanal monooxygenase (gene1604) participates in an intermediate step of alkane degradation, while acetaldehyde dehydrogenase (acetylating) (gene0790, gene2625, and gene2626, gene4367) facilitates aldehyde oxidation. Furthermore, acetyl-CoA C-acyltransferase (gene1860, gene4089, gene1401) represents a key enzyme involved in fatty acid β-oxidation.

These results suggest that petroleum degradation by strain B7-2 entails coordinated actions across multiple metabolic pathways, constituting an integrated degradation network. The aromatic hydrocarbon degradation pathway is characterized by the sequential activity of gene0804 (biphenyl-2,3-diol 1,2-dioxygenase), followed by gene4336/gene4337 (protocatechuate 3,4-dioxygenase), gene0791/gene2627/ gene4366 (4-hydroxy-2-oxovalerate aldolase), and gene0790/gene2625, among others (acetaldehyde dehydrogenase), ultimately leading to entry into the tricarboxylic acid (TCA) cycle. Similarly, the alkane degradation pathway involves gene1604 (alkanal monooxygenase), gene0138 (aldehyde dehydrogenase), and gene1860/gene4089 (acetyl-CoA C-acyltransferase), proceeding through fatty acid β-oxidation before integration into the TCA cycle.

### COG Functional Annotation

Protein sequences were functionally classified using the COG database within the eggNOG-mapper. As shown in Figure 5, 3,821 genes (78.19% of all predicted genes) were assigned to COG functional categories. The most abundant category was transcription (415 genes, 10.86%), followed by lipid transport and metabolism (360 genes, 9.42%), amino acid transport and metabolism (315 genes, 8.23%), and coenzyme transport and metabolism (289 genes, 7.56%). Additionally, a significant number of genes were associated with inorganic ion transport and metabolism (238 genes, 6.23%) and carbohydrate transport and metabolism (236 genes, 6.18%).

**FIG 5.**
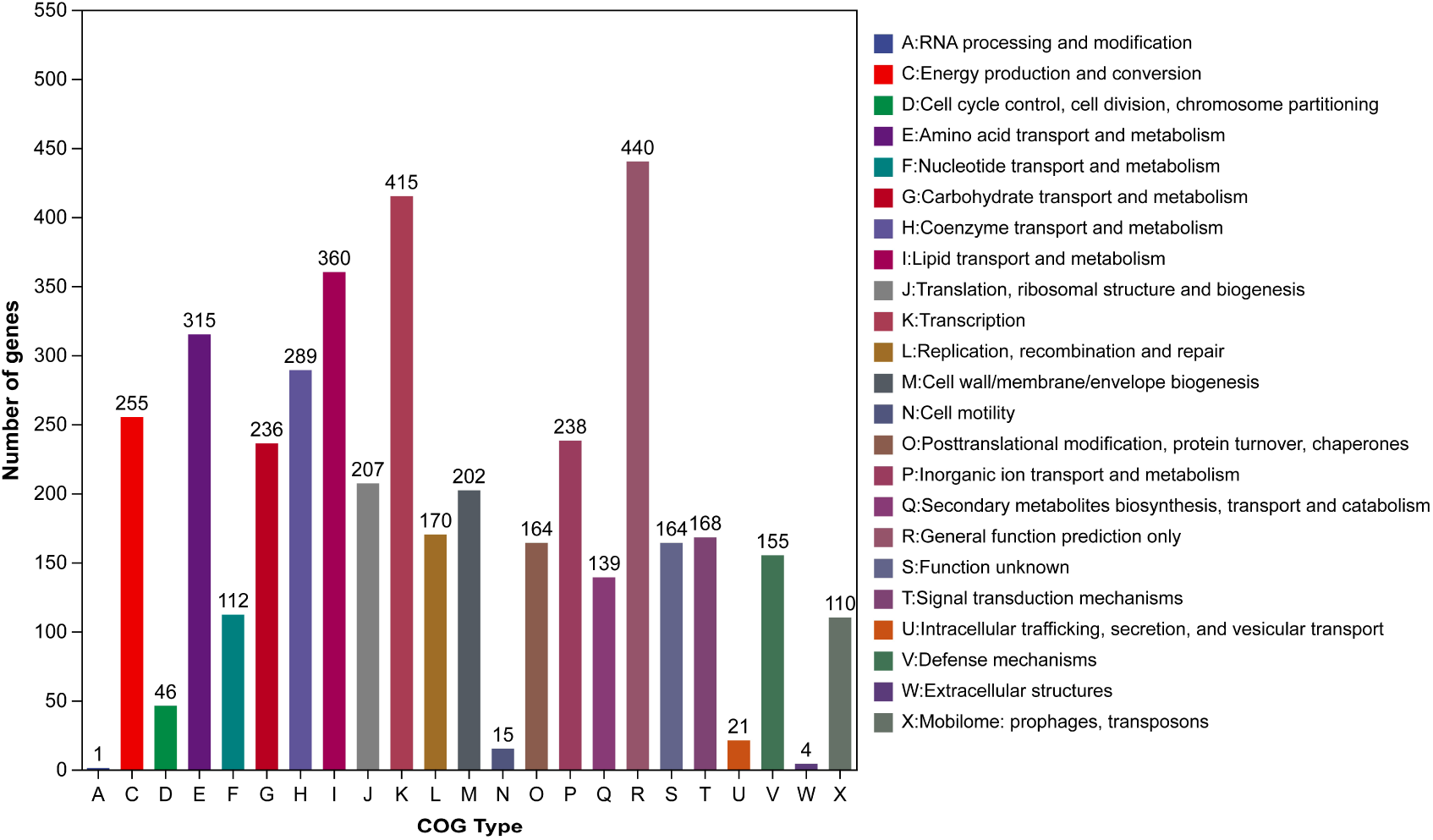
Functional categorization of predicted proteins in the *Gordonia* sp. B7-2 genome based on COG database.

### KEGG Annotation

Gene functions were annotated using the KEGG database. In total, 4,053 genes were assigned to KEGG pathways spanning six major functional categories: Metabolism, Cellular Processes, Genetic Information Processing, Organismal Systems, Human Diseases, and Environmental Information Processing, collectively encompassing 43 specific pathways (Figure 6). Among these categories, Metabolism was the most prominent, comprising 2,341 genes. Within metabolic pathways, 244 genes were associated with carbohydrate metabolism, accounting for 10.42% of metabolic genes, while 124 genes were associated with xenobiotic biodegradation and metabolism (5.30 %). A total of 225 genes were annotated for Environmental Information Processing and were primarily involved in membrane transport. Genetic Information Processing included 205 genes, whereas Cellular Processes and Human Diseases included 119 and 116 genes, respectively. The least represented category was Organismal Systems, with only 90 genes (0.04% of all annotated genes).

**FIG 6.**
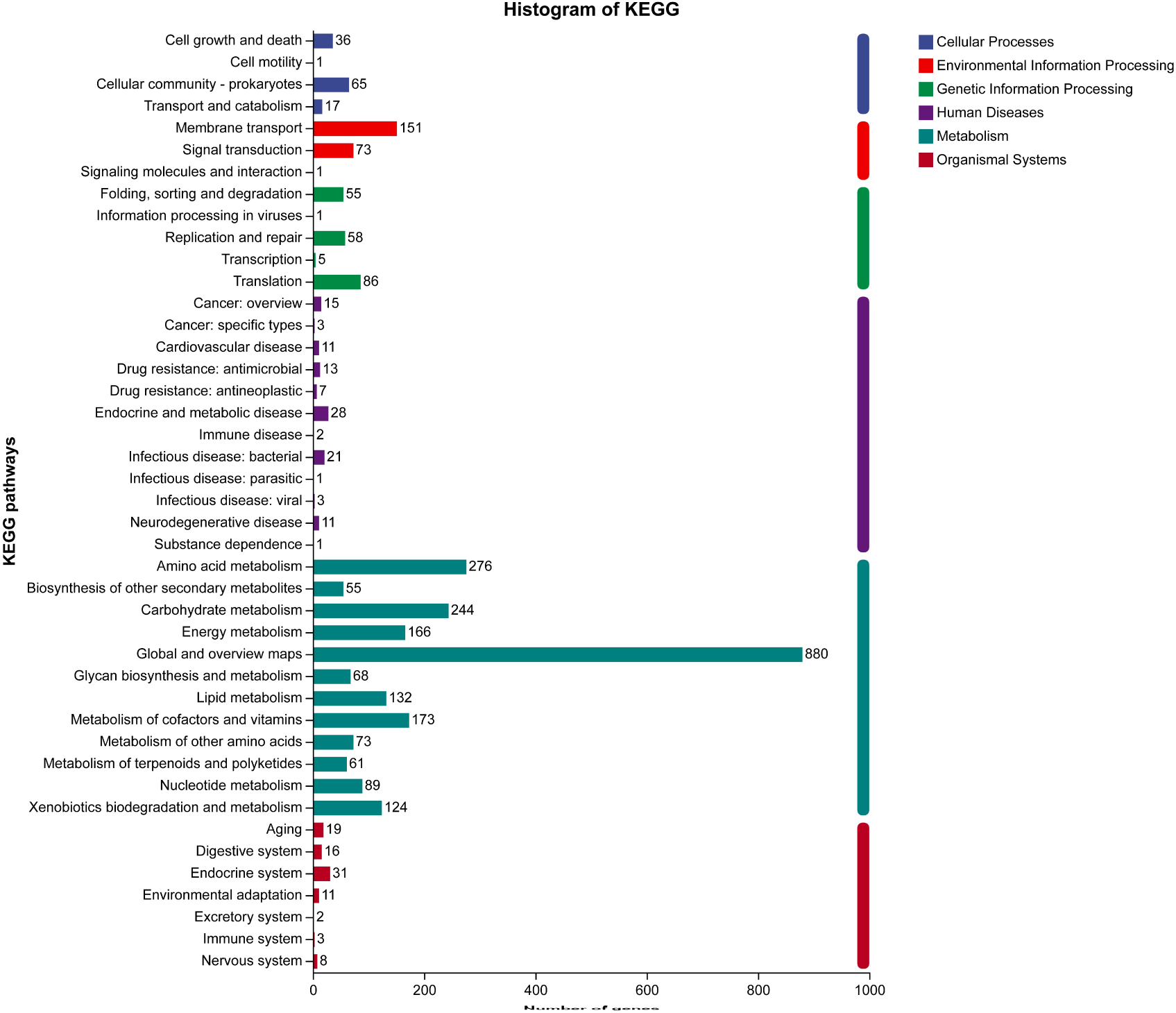
KEGG pathway assignment for the *Gordonia* sp. B7-2 genome.

Several key pathways associated with xenobiotic biodegradation were identified, including benzoate degradation (ko00362), xylene degradation (ko00622), chlorocyclohexane and chlorobenzene degradation (ko00625), styrene degradation (ko00643), and chloroalkane/chloroalkene degradation (ko00361) (Table 2). The presence of genes within these pathways indicates the potential of strain B7-2 to degrade petroleum-related pollutants.

**TABLE 2.** Key genes involved in xenobiotic biodegradation and metabolism pathways identified in the *Gordonia* sp. B7-2 genome.

| No. | Pathway ID | Description | Number of annotations |
| --- | --- | --- | --- |
| 1 | ko00623 | Toluene degradation | 2 |
| 2 | ko00622 | Xylene degradation | 18 |
| 3 | ko00626 | Naphthalene degradation | 3 |
| 4 | ko00624 | Polycyclic aromatic hydrocarbon degradation | 2 |
| 5 | ko00362 | Benzoate degradation | 56 |
| 6 | ko00642 | Ethylbenzene degradation | 1 |
| 7 | ko00643 | Styrene degradation | 7 |
| 8 | ko00361 | Chloroalkane and chloroalkene degradation | 6 |
| 9 | ko00625 | Chlorocyclohexane and chlorobenzene degradation | 12 |
| 10 | ko00071 | Fatty acid degradation | 60 |
| 11 | ko00980 | Metabolism of xenobiotics by cytochrome P450 | 3 |
| 12 | ko00903 | Limonene and pinene degradation | 23 |

Enzyme annotation revealed several key aromatic hydrocarbon-degrading enzymes, including initial dioxygenases such as naphthalene 1,2-dioxygenase (EC 1.14.12.12), catechol 1,2-dioxygenase (EC 1.13.11.1), protocatechuate 3,4-dioxygenase (EC 1.13.11.3), and benzoate 1,2-dioxygenase (EC 1.14.12.10), as well as monooxygenases including styrene monooxygenase (EC 1.14.15.2) and other related enzymes (EC 1.14.13.). Downstream enzymes, such as enoyl-CoA hydratase (EC 4.2.1.17) and 3-hydroxyacyl-CoA dehydrogenase (EC 1.1.1.35) were also identified, facilitating the channeling of aromatic ring cleavage products into central metabolic pathways. In contrast, key alkane-hydroxylating enzymes, including methane/alkane monooxygenases (EC 1.14.15.3), P450-type alkane monooxygenases (EC 1.14.14.), and non-heme iron monooxygenases (EC 1.14.13.), were not definitively annotated. The only annotated alkane 1-monooxygenase (EC 1.14.15.3) was associated with the naphthalene degradation pathway, while enzymes in limonene and pinene degradation primarily targeted cyclic olefins/terpenoids rather than linear or branched alkanes. Therefore, a complete alkane-degradation pathway was not identified in strain B7-2. These findings suggest that strain B7-2 possesses efficient aromatic hydrocarbon degradation capabilities, particularly via catechol and protocatechuate pathways for the metabolizing mono- and polycyclic aromatic hydrocarbons, including benzoate, toluene, xylene, ethylbenzene, naphthalene, and styrene.

### Annotation Based on NR and Swiss-Prot Databases

Functional annotation of predicted protein-coding genes was performed through sequence alignment against the non-redundant (NR) NCBI and Swiss-Prot databases. The NR database provides comprehensive sequence coverage and serves as a primary resource for gene annotation, whereas Swiss-Prot, maintained by the European Bioinformatics Institute, offers high-quality, manually reviewed annotations supported by experimental evidence. In total, 4,813 genes (98.46% of all predicted CDSs) were annotated using the NR database, and 3,400 proteins (69.57%) were assigned functional descriptions based on Swiss-Prot.

### Crude Oil Degradation Characteristics of Strain B7-2

The crude oil degradation capability of *Gordonia* sp. B7-2 was evaluated in minimal salts medium containing 300 mg/L crude oil as the sole carbon source. As shown in Figure 7, strain B7-2 degraded 35.98% of the crude oil within the first 7 days, reducing the concentration from 300 mg/L to 230 mg/L. Degradation efficiency increased to 42.41%, 60.08%, and 64.33% after 14, 21, and 28 days of incubation, respectively. The degradation process exhibited a strong positive correlation with incubation time (R² > 0.98). The average daily degradation rate was higher during the first 14 days (approximately 3.03%/day), followed by a decreased rate in the subsequent 14 days (approximately 1.57%/day). This pattern is consistent with typical microbial degradation kinetics, in which degradation rates decline as more readily degradable hydrocarbon fractions are consumed and metabolic adaptation progresses. The progressive accumulation of degradation over the 28-day period, reaching a final efficiency of 64.33%, indicates robust metabolic activity and effective microbial adaptation to crude oil as the sole carbon source.

**FIG 7.**
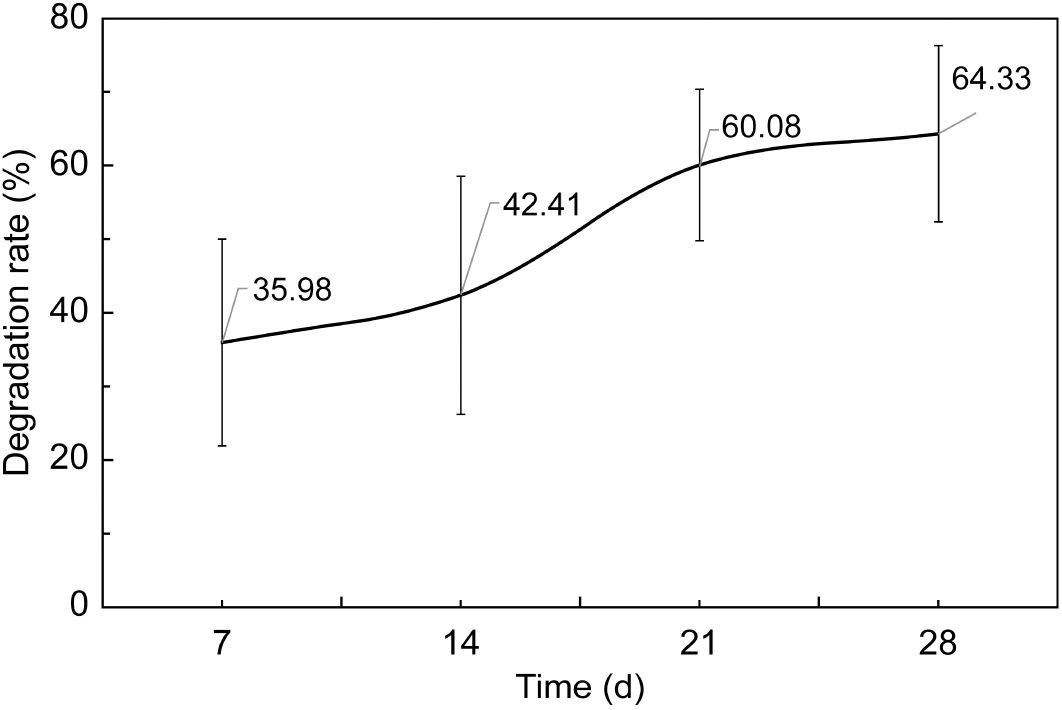
Crude oil degradation efficiency of *Gordonia* sp. B7-2 in minimal salts medium supplemented with 300 mg/L crude oil as the sole carbon source.

## DISCUSSION

The genus *Gordonia* has garnered considerable attention due to its capacity for hydrocarbon degradation and adaptation to contaminated environments. Nevertheless, the full genomic potential of many *Gordonia* strains, particularly those originating from underexplored ecological niches, remains insufficiently characterized. In this study, comprehensive genomic analysis of *Gordonia* sp. B7-2, isolated from mangrove sediments, elucidates the genetic basis underlying its specialized hydrocarbon degradation profile, thereby contributing to a deeper understanding of this biotechnologically important genus.

Polyphasic taxonomic analyses provided strong evidence supporting the taxonomic novelty of strain B7-2. The 16S rRNA gene sequence similarity values were below the recommended threshold for species demarcation (98.65%), and this observation was supplemented by ANI and dDDH values that were significantly lower than the established species boundaries (ANI<95–96% and dDDH <70%) (32, 33). These results support the classification of strain B7-2 as a putative new species. Recent phylogenetic studies have further indicated that the genus *Gordonia* comprises evolutionarily distinct lineages, raising the possibility of taxonomic refinement at the genus level. The isolation of strain B7-2 from mangrove sediments, a unique and biologically rich ecosystem, highlights the importance of exploring understudied habitats to for uncover microbial diversity with specialized functional traits (34). Mangrove ecosystems, in particular, have emerged as reservoirs of *Gordonia* species with unique metabolic capabilities, as evidenced by the identification of *Gordonia hongkongensis* from octocoral (35). Moreover, comprehensive genomic analyses of *Gordonia* species have revealed extensive catabolic versatility, supporting the notion that such environments may harbor novel *Gordonia* lineages with specialized metabolic functions for environmental adaptation and bioremediation applications (36). The development of genus-specific primers has further facilitated the detection of previously unrecognized *Gordonia* species from diverse environments, reinforcing the phylogenetic diversity of this genus and the importance of continued exploration of specialized ecological niches, including mangrove sediments.

Genome annotation provided insights into the genetic determinants underlying the hydrocarbon degradation capabilities of strain B7-2. Notably, a complete set of genes associated with aromatic hydrocarbon degradation was identified, including those encoding key enzymes such as protocatechuate 3,4-dioxygenase (PCD) and biphenyl-2,3-diol 1,2-dioxygenase (BPDO). PCD, an intradiol dioxygenase, catalyzes the critical ring cleavage of 3,4-dihydroxybenzoate within the β-ketoadipate pathway, whereas BPDO, an extradiol dioxygenase, acts on biphenyl-2,3-diol in the meta-cleavage pathway (37, 38). These pathways are fundamental for the breakdown of mono- and PAHs (38, 39). The genomic features of strain B7-2 are broadly consistent with those of other *Gordonia* strains, such as *G. polyisoprenivorans* 135, whose complete genome also reveals a rich repertoire of PAH catabolic genes, albeit with variations in gene organization that may influence degradation efficiency and flexibility (40). The predicted genomic potential of strain B7-2 was phenotypically corroborated by the efficient degradation of crude oil, which achieved 64.33% removal over 28 days. The degradation kinetics, characterized by a higher daily degradation rate during the initial incubation period followed by a gradual decline, are consistent with first-order degradation models commonly reported for the aromatic hydrocarbon metabolism by *Gordonia* species (41). An additional notable genomic feature is the absence of a complete pathway for the initial oxidation of linear alkanes, indicating a metabolic preference toward aromatic hydrocarbons. This specialization suggests that strain B7-2 may fulfill a complementary ecological role within microbial consortia, where cooperation with alkane-degrading microorganisms enables more comprehensive crude oil degradation through metabolic cross-feeding (39).

Beyond its hydrocarbon-degrading capacity, the genome of B7-2 harbors a diverse array of BGCs, as predicted by antiSMASH 6.0 (31), particularly those encoding non-ribosomal peptide synthetases and polyketide synthases (42). This finding highlights the largely unexplored potential of this strain to produce novel secondary metabolites, a trait increasingly recognized within the genus *Gordonia* for yielding compounds with diverse chemical structures and bioactivities (43). The genomic capacity for natural product biosynthesis suggests that the ecological role of strain B7-2 extends beyond hydrocarbon degradation, and potentially involves the production of siderophores for iron acquisition or antimicrobials for niche competition (44). In addition, annotation of carbohydrate-active enzyme (CAZyme) genes indicates a robust ability to degrade plant polysaccharides, potentially facilitating beneficial interactions with plants in the mangrove rhizosphere. This trait could support nutrient acquisition from root exudates and decaying plant matter and suggests a synergistic role for strain B7-2 as a plant growth-promoting bacterium within its native habitat. The coexistence of specialized catabolic and biosynthetic capabilities in strain B7-2 highlights the metabolic versatility of the genus *Gordonia*, supporting its ability to thrive in complex environments such as mangrove sediments through diverse microbial interactions and nutrient cycling.

The methodological framework employed in this study, integrating complete genome sequencing with comparative genomic analysis, enabled a functional and ecological interpretation of strain B7-2 beyond a descriptive gene inventory. Degradative and biosynthetic gene clusters were systematically identified using genome mining tools, including antiSMASH 6.0 (31). Comparative genomic analysis with other hydrocarbon-degrading bacteria further revealed the distinctiveness of the B7-2 genome, which encodes a specialized aromatic degradation apparatus while lacking key pathways for alkane oxidation. This genomic configuration suggests that strain B7-2 occupies a relatively narrow metabolic niche (45). By providing a predictive blueprint For metabolic capability, this genome-guided approach helps explain the observed degradation kinetics and supports the view that strain B7-2 can function as a specialized member of synthetic microbial consortia, targeting aromatic hydrocarbon fractions in partnership with alkane-degrading microorganisms. In this context, the genome serves both as a record of environmental adaptation and as a rational design manual for potential biotechnological applications, demonstrating how integrated genomic mining can extract actionable insights from sequence data.

While the integrated genomic and phenotypic approach provides a robust blueprint of the metabolic capabilities of strain B7-2, several limitations of the present study should be noted to guide future research. First, the catabolic predictions are based on genomic annotation and homology, and would benefit from direct validation through transcriptomic or proteomic analyses to confirm the expression and regulation of key degradation genes under induction conditions. Second, the degradation efficiency and kinetics were characterized in controlled laboratory-scale assays. The performance and ecological relevance of the strain should be further evaluated under more complex in situ or simulated environmental conditions, which involve factors such as nutrient competition, fluctuating physicochemical parameters, and native microbial communities. Finally, the analysis measured total crude oil removal without distinguishing specific hydrocarbon fractions. Future work employing fraction-specific or compound-level analysis would provide finer resolution on the substrate preferences of strain B7-2 and the recalcitrance of particular oil components. Addressing these aspects will be crucial for translating the promising genomic potential of strain B7-2 into effective and reliable bioremediation applications.

## CONCLUSION

The present study provides genomic and phenotypic characterization of *Gordonia* sp. B7-2, a petroleum-degrading bacterium isolated from mangrove sediments. Analysis of its complete genome sequence revealed a specialized catabolic apparatus encoding a complete aromatic hydrocarbon degradation pathway, while genes associated with the initial oxidation of alkanes were absent. Phenotypic validation confirmed this functional specialization, as strain B7-2 degraded 64.33% of crude oil within 28 days, with higher degradation rates observed during the initial 2 weeks than in the subsequent period. Together, these findings highlight a strategy of metabolic specialization towards aromatic compounds and demonstrate how genomic insights can predict and explain functional traits in hydrocarbon-degrading bacteria. This study expands the genomic resources available for the biotechnologically valuable genus *Gordonia* and provides a rational basis for designing specialized microbial consortia targeting aromatic contaminants for bioremediation applications.

## AUTHOR CONTRIBUTIONS

Conceptualization: F.J., H.F., and X.X.; Methodology: H.S., H.F., and X.X.; Software: X.X., F.J., and H.S.; Investigation: H.S., Z.Z., and M.L.; Data Curation: X.X. and H.F.; Writing — Original Draft Preparation: F.J., H.S., and H.F.; Writing—review and editing, Z.Z., M.L., F.J., X.X., and H.F.; Project Administration: X.X. and H.F. All authors have read and agreed to the published version of the manuscript.

## ACKNOWLEDGEMENTS

This study was supported by the natural science fund projects of Hainan Province (Grant No. 424RC525); the Scientific Research Foundation of Hainan Tropical Ocean University (Grant No. RHDRC202338); the Key Research Program of Qiongtai Normal University (Grant No. qtky202401); the Scientific Research Foundation of Qiongtai Normal University (Grant No. 30030007040); and Hainan Province College Students’ Innovation and Entrepreneurship Program (Grant No. S202511100015).

## DATA AVAILABILITY

The complete genome sequence has been deposited in the NCBI database under accession number CP194566 (https://www.ncbi.nlm.nih.gov/nuccore/CP194566.1/).

---

**Table S1.** General genomic features of *Gordonia* sp. B7-2.

| Item | Number | Item | Number |
| --- | --- | --- | --- |
| Genome size (bp) | 5,393,941 | Number of genomic islands | 20 |
| Genome GC content (%) | 65.99 | Total length of gene island (bp) | 507,403 |
| Number of genes | 4,887 | Number of prophages | 2 |
| Total length of gene (bp) | 4,879,074 | Total length of prophages (bp) | 29,918 |
| Total length of coding region (%) | 90.45 | Number of tandem repeats | 218 |
| Average length of gene (bp) | 998.38 | Number of short interspersed elements | 11 |
| Gene Density (kb) | 0.91 | Number of long interspersed repeated sequences | 5 |
| Number of rRNA genes | 6 | Number of Transposon | 20 |
| Number of tRNA genes | 46 | Number of Insertion Sequence | 24 |
| Number of sRNA genes | 39 | Total potential CRISPRs | 30 |

